# Prior use-dependent plasticity triggers different individual corticomotor responses during persistent musculoskeletal pain

**DOI:** 10.1101/2025.01.15.633250

**Authors:** Anna M. Zamorano, Boris Kleber, Enrico De Martino, Ainhoa Insausti-Delgado, Peter Vuust, Herta Flor, Thomas Graven-Nielsen

**Author notes:** **Corresponding author:** Anna M. Zamorano, Ph.D., Center for Music in the Brain (MIB)., Department of Clinical Medicine, Aarhus University., Building 1710, Universitetsbyen 3. 8000 Aarhus C, Denmark.

## Abstract

Movement repetition is crucial for pain interventions. It facilitates the rehabilitation of motor patterns, the acquisition of motor skills and the genesis of adaptive use-dependent plasticity. However, the influence of prior motor experience and pre-existing use-dependent plasticity on pain severity and progression remains poorly investigated. This study investigated the effects of pre-existing use-dependent plasticity during the development of prolonged experimental musculoskeletal pain. Using transcranial magnetic stimulation, corticospinal excitability was assessed by measuring the rest-motor thresholds (RMTs), motor-evoked potential (MEP), representational area of the motor map, volume, and center of gravity of the first dorsal interosseous (FDI) muscle in musicians (n=19), a well-known ecological model of use-dependent plasticity, and in non-musicians (n=20). All participants attended three sessions (Day1, Day3, Day8). Prolonged pain for several days was induced by intramuscular injection of nerve growth factor (NGF) into the right FDI muscle at the end of Day1. Compared to Day1, prolonged pain uniquely led to reduced motor map volume in non-musicians on Day3 (*p*=0.004), who also showed higher NGF-related pain intensity compared to musicians. The motor maps of musicians, which were already smaller in pain-free conditions (Day1) compared to non-musicians (*p*=0.021), remained non-significantly different across days. Notably, corticomotor responses (map volume, MEP amplitude, and RMTs) at Day1 were correlated to weekly and accumulated musical training. These findings demonstrate that pre-existing use-dependent plasticity associated with motor training may counteract the effects of prolonged pain in the motor system. Moreover, it confirms that prior motor experience acts as a source of individual variability to pain.

## INTRODUCTION

When we adapt to changing environmental conditions and experiences, our motor system undergoes plastic changes. Pain, in particular, is a potent stimulus capable of shaping our behaviour and modulating our motor pathways [9,45,46]. This happens, for example, when acute pain triggers withdrawal reflexes as a protective measure to prevent further harm [41]. The adaptive response is accompanied by impaired movement control and a temporal reduction in corticomotor excitability (CE), a key measure of the brain’s responsiveness to motor commands [4,9,16,35]. Chronic pain, on the other hand, is also associated with long-lasting functional reorganization of the primary motor cortex (M1) [37,39]. However, corticomotor responses are highly variable during persistent pain conditions [6,40,43]. Results from recent meta-analysis and machine learning studies investigating these motor adaptations have shed light on this variability, indicating that reduced corticomotor maps are closely linked to increased pain severity during sustained pain [8,9]. Nevertheless, the underlying mechanisms driving this variability remain unknown.

Use-dependent plasticity in the nervous system occurs as a result of repeated stimulation and experience [5,29]. When we repeat a motor task to learn a new skill for hours to weeks, an enhancement of corticomotor excitability occurs in the M1 [10,27,31]. This process is mediated by various mechanisms, such as long-term potentiation (LTP) [3,5,15,27,34]. When this training continues over time (months to years), extensive sensorimotor training leads to reduced CE and refined functional motor maps [22,26,30,47]. Those changes are indicative of overlearning and automaticity of tasks, such as the fine-motor skills required for musical practice [18]. While short-term motor training has been investigated to understand its effects on pain processing, the interaction of long-term use-dependent plasticity with pain development is not fully understood.

Musicians are a unique population to study the impact of long-term sensorimotor training in sensory processing [13,28,32,33]. In pain processing, recent studies with musicians highlight that prior use-dependent plasticity may lead to enhanced phasic pain sensitivity and increased cortical-evoked responses to painful stimulations in pain-free individuals [49,51]. Notably, when musicians have chronic pain, this enhanced phasic pain sensitivity is still present [48], but their motor responses are not affected compared to pain-free musicians [47]. Moreover, they report fewer interferences with daily activities and reduced insula-based connectivity with pain-related brain regions [50]. This suggests that the long-term use-dependent plasticity contributes to inter-individual variability in pain processing.

This experiment aimed to study the interaction between long-term use-dependent plasticity and prolonged muscle pain development. To this, we injected nerve growth factor (NGF) into a hand muscle in musicians and non-musicians, a procedure which allows for studying the motor system changes as pain develops during up to 14 days [1,38]. Using transcranial magnetic stimulation (TMS), the corticomotor excitability was assessed by exploring the motor-evoked potentials (MEPs), map area, map volume, and centre of gravity of the first dorsal interosseous (FDI) muscle during the development of prolonged muscle pain in non-musicians and musicians. We hypothesised that musicians’ corticomotor maps would remain stable during the development of prolonged muscle pain, while, in non-musicians, they would decrease. We further anticipated that corticomotor depression would be associated with worse pain and disability and worse interferences with daily activities.

## MATERIALS AND METHODS

### Participants

Thirty-nine participants were recruited via calls mainly at Aalborg University, Aarhus University, and The Royal Academy of Music, Aarhus/Aalborg. Nineteen of these were healthy musicians (6 females, mean age 25.0 ± 2.6 years) consisting of 9 amateurs (4,406 ± 2,776 hours of accumulated training and 8.1 ± 3.7 hours of weekly training) and 10 conservatory-trained instrumentalists (15,540 ± 6,621 hours of accumulated training and 27.6 ± 14.2 hours of weekly training). The control group included 20 healthy non-musicians (nine female, 19 right-handed, mean age 26.9 ± 5.3 years) without any prior formal or informal music training. Both groups participated in a previous study reporting psychological measures (State-Trait Anxiety, pain catastrophizing, and pain vigilance), nociceptive and non-nociceptive-evoked potentials, NGF-pain distribution and ratings, as well as quantitative sensory assessment (pressure pain thresholds, and electrical perception thresholds) [48]. Exclusion criteria were neurological, cardiorespiratory, or mental disorders, or pregnancy, as well as history of chronic pain or current acute pain. Participants completed a transcranial magnetic stimulation (TMS) safety screen[20] prior to commencement. The sample size was estimated using G*Power[14] and based on previous publications using a similar approach[12,38,47] to ensure 80% power for detecting at least a medium effect size (Cohen’s d ≥ 0.6) on motor maps with a repeated measures analysis of variance (ANOVA) at an alpha level of 0.05. The sample size required for each primary outcome (nociceptive evoked potentials, published elsewhere, and corticomotor maps, current publication) was calculated separately, and the larger of the two sample sizes was chosen to ensure sufficient power for both outcomes. All participants received written and verbal information about the study and provided written consent. The study was registered on ClinicalTrials.gov (NCT04457466) and performed in accordance with the Declaration of Helsinki[2], approved by the local ethics committee (Den Videnskabsetiske Komité for Region Nordjylland, N-20170040).

### Experimental Procedure

The experiment involved 3 sessions (Day1, Day3, and Day8) over 8 days. Participants were seated in a comfortable chair for the laboratory sessions. At the end of Day1, all participants received an injection of NGF into the right first dorsal interosseous (FDI) muscle to induce prolonged muscle pain in the hand for several days. At the beginning of each session, participants reported the demographic (Day1) and the pain-related questionnaires (Day1, Day3, Day8). Subsequently, on Day1 (before the NGF injection), Day3, and Day8, corticomotor excitability (MEPs) and motor maps were assessed by using TMS over the cortical representation of the FDI muscle).

### Experimental Prolonged Muscle Pain

Muscle pain and hyperalgesia were induced by intramuscular injections of NGF into the right FDI muscle after all assessments on Day1. Sterile solutions of recombinant human beta-NGF were prepared by the pharmacy (Skanderborg Apotek, Denmark). The site of injection was cleaned with alcohol, and NGF solution (5μg/0.5 mL) was immediately injected into the muscles of the right hand. Pain ratings in resting conditions were reported at the beginning of Day3 and Day8 using a Numeric Rating Scale (NRS) with 0 defined as “no pain” to 10 corresponding to “worst pain imaginable”. Pain pressure thresholds (PPTs) were assessed at each session (Day1, Day3, and Day8) using a handheld pressure algometer (1-cm2 probe, Somedic Electronics, Solna, Sweden) covered by a disposable latex sheath. Pressure was applied perpendicularly to the skin over the left and right FDI muscle until participants reported pain. The PPTs and NRS pain ratings reported in this study are described in detail in Zamorano et al. [48].

### Disability of Arm, Shoulder, and Hand questionnaire

The Disabilities of the Arm, Shoulder, and Hand (DASH) questionnaire was used to evaluate the hand function and symptoms before and during the experimentally induced hand pain [19]. The DASH self-assessment questionnaire contains 30 items, plus 2 optional modules of 4 items (sports/performing arts or work) that have the goal to identify the specific difficulties that professional athletes/performing artists or other groups of workers might experience.

### Corticomotor Excitability

Electromyographic (EMG) activity was recorded and pre-processed following the same procedure as in Zamorano et al. [47]. EMG activity was obtained from the muscle belly of the right FDI using silver/silver chloride surface electrodes (Neuroline 720-01-K, Ambu® A/S). The EMG signals were sampled at 4 kHz, pre-amplified (1000 × gain), and bandpass filtered between 5 Hz and 2 kHz. The data were digitized by a 16-bit data acquisition card (National Instruments, NI6122) and saved using custom-made Labview software (Mr. Kick, Knud Larsen, SMI, Aalborg University). EMG activity was pre-processed offline using Matlab (The Math-Works, Natick, USA). First, the TMS stimulation artefact was removed and the voltage difference at the extremities of the segments was corrected before being merged. The resulting signal was bandpass filtered between 5 and 1000 Hz and notch filtered (50 Hz) using a 2nd-order Butterworth filter.

Single-pulse TMS were delivered using a Magstim 200 stimulator (Magstim Co. LtD) and a figure-of-eight coil. The coil was positioned over the left hemisphere at a 45-degree angle to the sagittal plane to preferentially induce current in a posterior-to-anterior direction. All TMS procedures adhered to the TMS checklist for methodological quality [7]. Participants were seated comfortably in a chair and familiarised with the transcranial magnetic stimulation before starting the experiment. First, the optimal cortical site (hotspot) of the right FDI muscle, determined as the coil position that evoked a maximal peak-to-peak MEP for a given stimulation intensity, was obtained. Then, the resting motor threshold (rMT), defined as the minimum stimulator intensity at which 5 out of 10 stimuli applied at the optimal scalp site evoked a response with a peak-to-peak amplitude of at least 50 μV [36]. Finally, ten MEPs were recorded at 120% of rMT over the optimal cortical site with the FDI muscle at rest to evaluate corticomotor excitability. MEP responses were measured as peak-to-peak amplitudes and averaged for analysis.

### Motor Cortical Maps

The procedure for cortical mapping was done following previous studies [12,47]. Participants were fitted with a cap, marked with a 1 × 1 cm grid and centred on the vertex (point 0,0). The stimulus intensity for mapping was 120% rMT. TMS was applied every 6 s with a total of 5 stimuli at each site. The scalp sites were pseudo-randomly measured from the hotspot until no MEP was detected (defined as <50 µV peak-to-peak amplitude in all 5 trials in all border sites. Trials containing background EMG activity were discarded from the analysis.

The number of active grid sites and map volume were calculated. A grid site was considered “active” if the mean peak-to-peak amplitude of the 5 MEPs evoked at that site was greater than 50 µV. The mean peak-to-peak MEP amplitudes in µV at all active sites were summed to calculate the map volume. The map area was calculated by summing the number of active sites. The centre of gravity (CoG) was defined as the amplitude-weighted centre of the map and was calculated for each muscle using the formula:

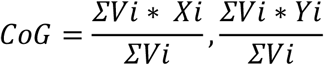

where Vi represents mean MEP amplitude at each site with the coordinates Xi, Yi.

### Statistical Analysis

Data are presented as means and standard deviation in text, tables, and figures. Data were statistically analysed with IBM SPSS Statistics 29 for Windows. Data were screened for assumptions of normality, sphericity, homogeneity, and independent errors using descriptive plots and statistical tests. Corticomotor data (RMTs, MEPs, map area, map volume, and number of discrete peaks) were compared across *Time* (Day1, Day3, and Day8, and *Group* (musicians vs non-musicians; between group factor) using two-way repeated-measures ANOVA. Significant main factors or interactions were analysed post hoc using Bonferroni’s procedure to correct for multiple comparisons.

Correlations were computed to investigate whether the corticomotor responses (rMTs, MEPs, map area, and map volume) during pain-free conditions (Day1) and the peak of NGF pain effects (Day3) could be associated with pain perception and disability (PPTs, pain NRS ratings and DASH scores) on Day3, across all participants. In musicians, it was furthermore tested if the accumulated and weekly training affected the corticomotor data in pain-free conditions (Day1). For all tests used, the level for statistically significant difference was set at *p* < 0.05. Bonferroni’s procedure was used to correct for multiple comparisons or correlations.

## RESULTS

One participant had a family history of epilepsy and was excluded from the TMS assessments. Results about distribution and intensity of NGF-induced muscle pain, as well as PPTs are described in Zamorano et al.[48]. In summary, NGF-experimental muscle pain distribution was comparable in both groups of participants, affecting the local dorsal and palmar regions of the hand on Day3 and Day8. PPTs were reduced in both groups and both right and left hands (i.e., local injection site and contralateral hand) on Day3 and Day8, not showing significant group differences. However, NRS scores of NGF-induced pain were higher in non-musicians than in musicians.

### Corticomotor Excitability

The resting motor threshold was not significantly different between groups and across days (Table 1; *p* < 0.633 and *ƞ*^*2*^ < 0.01).

**Table 1.** Corticomotor excitability measures and psychometric scores (mean ± SD) for musicians and non-musicians.

|  | Musicians (n = 19) |  |  | Non-musicians (n = 19) |  |  |
| --- | --- | --- | --- | --- | --- | --- |
|  | Day1 | Day3 | Day8 | Day1 | Day3 | Day8 |
| **Corticomotor excitability** |  |  |  |  |  |  |
| rMT (%) | 38 ± 6 | 38 ± 5 | 38 ± 6 | 40 ± 7 | 41 ± 6 | 40 ± 5 |
| CoG lat (cm) | 0.9 ± 0.8 | 1.1 ± 0.6 | 1.0 ± 0.5 | 1.2 ± 0.7 | 1.3 ± 0.6 | 1.0 ± 0.6 |
| CoG long (cm) | 4.7 ± 0.7 | 4.8 ± 0.8 | 4.8 ± 0.8 | 5.1 ± 0.7 | 5.0 ± 0.6 | 4.8 ± 0.8 |
| Map volume (mV) | 5196 ± 3611* | 6504 ± 5185 | 5917 ± 4818 | 9353 ± 6545* | 5410 ± 4361# | 8090 ± 5967 |
| Map area (n) | 14.1 ± 5.7Φ | 15.7 ± 5.6Φ | 14.8 ± 4.6Φ | 19.6 ± 9.0Φ | 16.7 ± 6.0Φ | 19.8 ± 8.2Φ |
| MEP (μV) | 847 ± 569 | 1103 ± 930 | 990 ± 729 | 1273 ± 730 | 844 ± 624 | 1172 ± 1016 |
| Active sites (n) | 1.6 ± 0.7Φ | 1.5 ± 0.8Φ | 1.3 ± 0.6Φ | 1.7 ± 0.6Φ | 1.9 ± 0.9Φ | 2.1 ± 1.1Φ |
| **Psychometrics** |  |  |  |  |  |  |
| DASH | 2.2 ± 2.8 | 13.3 ± 9.5 | 9.8 ± 6.7 | 2.9 ± 4.5 | 13.4 ± 11.7 | 12.0 ± 11.6 |
| DASH music | 1.6 ± 3.6 | 19.1 ± 19.7 | 12.2 ± 12.9 | NA | NA | NA |

rMT: rest motor threshold; MEP: motor evoked potential; CoG: centre of gravity; DASH: Disabilities of the Arm, Shoulder and Hand; NA: not applicable. Significantly different between groups on the day (\*, $P < 0.05$ ). Significantly different compared to Day1 and Day8 within the group (#, $p < 0.05$ ). Significantly different between groups ( $\Phi$ , $p < 0.05$ ).

The ANOVA of the maximum MEP showed a significant *Group x Time* interaction (*F*_1, 72_ = 3.573; *p* = 0.033; *ƞ*^*2*^ = 0.09). Post-hoc analysis indicated a trend towards a significantly lower MEP in musicians compared with non-musicians on Day1 (*p* = 0.052) and reduced MEP Day3 compared with Day1 in non-musicians (*p* = 0.056). No significant *Time* effect (*F*_1, 72_ = 0.382; *p* = .684; *ƞ*^*2*^ < 0.01) or *Group* effect (*F*_1, 36_ = .324; *p* = .573; *ƞ*^*2*^ < 0.01) were found.

### Corticomotor Maps

The ANOVA of the map volume (Fig. 1A and 1B, Table 1) showed a significant *Group x Time* interaction (*F*_2, 72_ = 5.320; *p* = 0.007; *ƞ*^*2*^ = 0.13). Post-hoc analysis indicated that the map volume of musicians and non-musicians was different on Day1 (*p* = 0.021). Moreover, non-musicians showed a reduction in map volume on Day3 compared to Day1 (*p* = 0.004) and Day8 (*p* = 0.022). No significant *Time* effect (*F*_2, 72_ = 1.465; *p* = 0.238; *ƞ*^*2*^ = 0.11) or *Group* effect (*F*_2, 36_ = 1.572; *p* = 0.218; *ƞ*^*2*^ = 0.04) were found.

**Figure 1.**
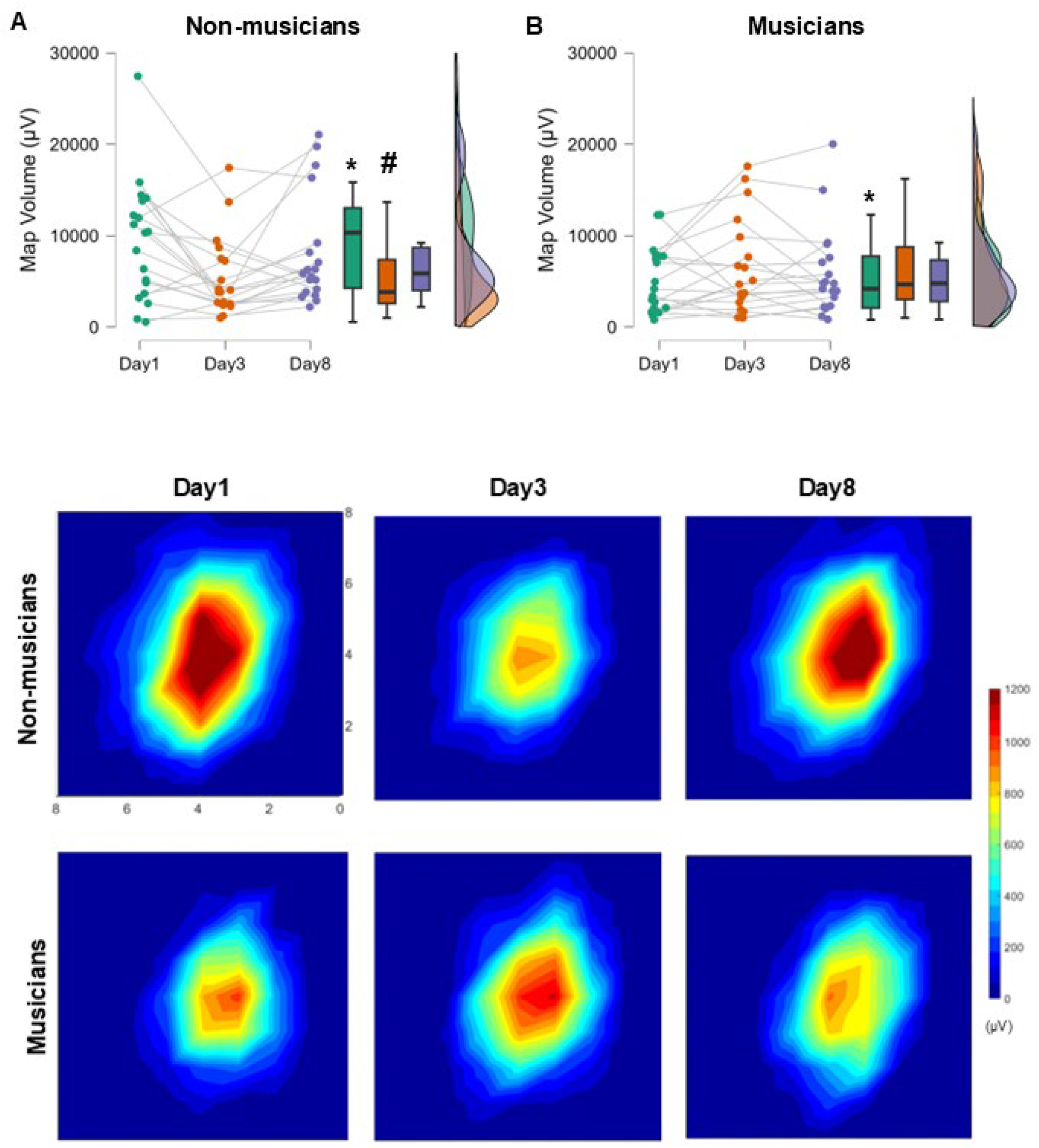
Corticomotor measures and maps in musicians and non-musicians. Changes in motor map volume (μV) across time in non-musicians **(A)** and musicians **(B)**. Individual dots represent the mean values for each participant at Day1 (green), Day3 (orange), and Day8 (purple). Boxplots summarize group-level data, showing the median, interquartile range, and overall variability. Violin plots on the right illustrate the distribution of map volumes across all time points for each group. **C)** Map area representation obtained for the right FDI muscle across days in musicians and non-musicians. Axes coordinates are referenced to the stimulation site that evoked the greatest motor-evoked potential (0,0 is the centre grid reference in the map) obtained for each individual. The X-axis represents the posterior side of the motor cortex, while the Y-axis represents the medial side of the motor cortex. Significantly different between groups on the day (*, P < 0.05). Significantly different compared to Day1 and Day8 within the group (#, P < 0.05).

The ANOVA of the map area (Fig. 1C, Table 1) showed a significant *Group* effect (*F*_2, 36_ = 4.504; *p* = 0.041; *ƞ*^*2*^ = 0.11), where musicians showed reduced motor map area compared to non- musicians. A trend approaching significance was found for the *Group x Time* interaction (*F*_2, 72_ = 2.801; *p* = 0.067; *ƞ*^*2*^ = 0.07). No significant *Time* effect (*F*_2, 72_ = 0.579; *p* = 0.563; *ƞ*^*2*^ < 0.02) was found for the map area.

The ANOVA of the number of active grid sites (Table 1) showed a significant *Group* effect (*F*_1, 36_ = 5.053; *p* = 0.020; *ƞ*^*2*^ = 0.14), with musicians having a lower number of active sites compared to non-musicians. No significant *Time* effect (*F*_1.65, 72_ = 0.157; *p* = 0.855; *ƞ*^*2*^ < 0.01) or interaction *Time x Group* (*F*_1.65, 72_ = 2.248; *p* = .113; *ƞ*^*2*^ = 0.06) were found.

The mediolateral center-of-gravity (CoG) location of the motor map was not significantly different between groups or across days (Table 1; all *p* < 0.157 and *ƞ*^*2*^ < 0.05).

### No association between corticomotor responses with pain ratings and pain disability

Among all participants, the corticomotor responses on Day1 or Day3 were not significantly associated with NRS pain ratings and DASH scores on Day3 (all *p* > 0.05).

### Higher rest motor thresholds are associated with increased pressure pain perception

Among all participants, rest motor thresholds on Day1 and Day3 (Fig 2A) correlated negatively with pressure pain thresholds on Day3 (r = -0.35, p = 0.029 and r = -0.33 p = 0.04, respectively), indicating that those participants with higher rest motor thresholds showed higher pressure pain perception during the transition to persistent pain.

**Figure 2.**
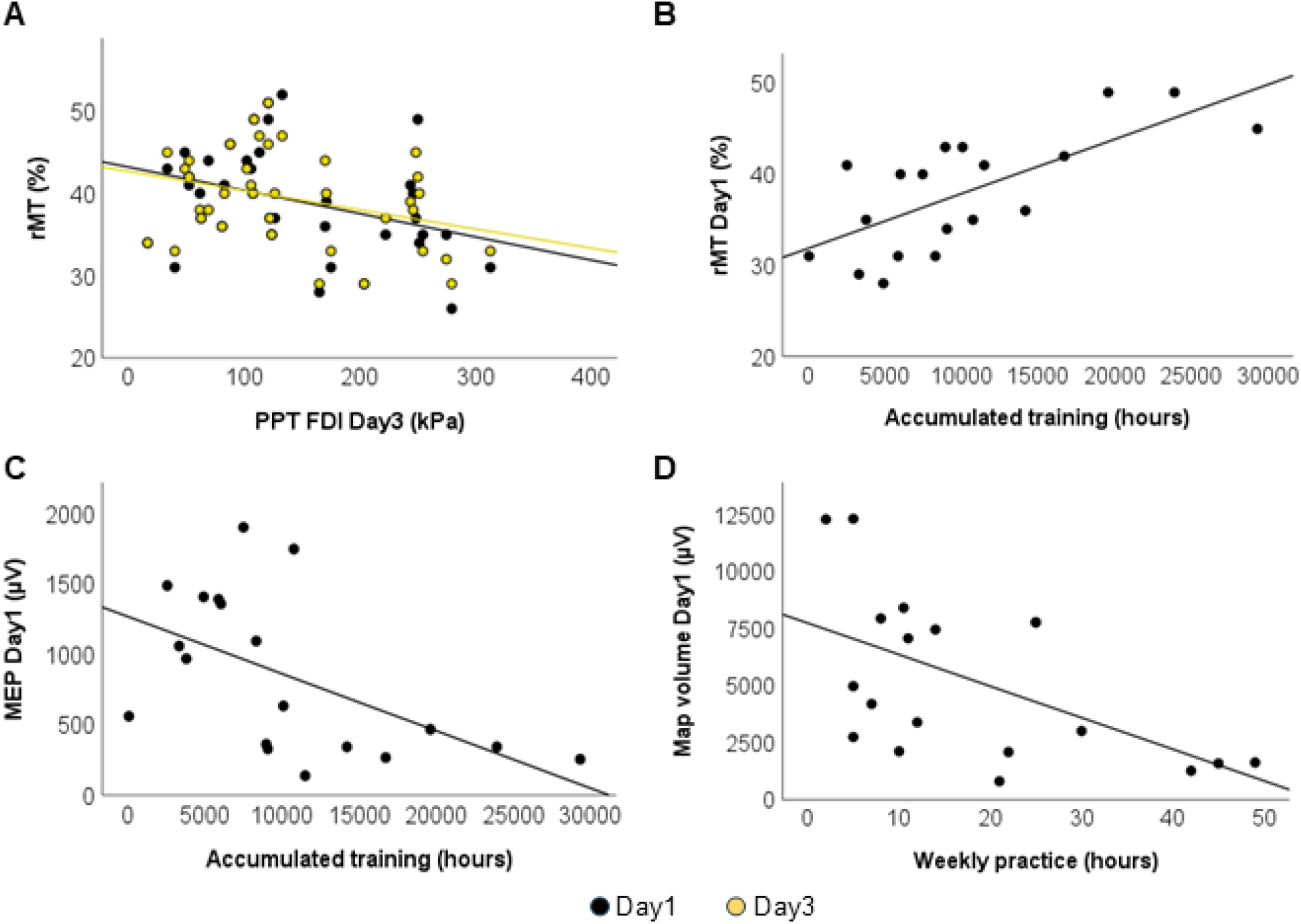
Significant correlations of corticomotor outputs, pain sensitivity, and musical practice. **A)** Among all participants, rest motor thresholds on Day1 (black dots) and Day3 (yellow dots) correlated negatively with pressure pain thresholds on Day3. **B)** In musicians, the accumulated musical training positively correlated with the RMT on Day1. **C)** The maximum MEP amplitude on Day1 inversely correlated with the accumulated musical training. **D)** The weekly musical training inversely correlated with the map volume on Day1.

### In musicians, the weekly and accumulated musical training is associated with the map volume, the MEP amplitude, and the rest motor threshold

In musicians, the accumulated musical training was correlated with the rMTs on Day1 (Fig. 2B, *r* = 0.70, *p* < 0.001) and negatively correlated with the MEP amplitude on Day1 (Fig. 2C, *r* = -0.54, *p* = 0.017). Moreover, the weekly training was negatively correlated with the map volume on Day1 (Fig. 2D, *r* = -0.55, *p* = 0.015) and with the MEP amplitude on Day1 (*r* = -0.56, *p* = 0.013).

## DISCUSSION

Here, we found that prolonged pain uniquely led to a reduction of map volume in non-musicians, indicating a reduction in corticomotor excitability as a function of prolonged muscle pain after injection of NGF. Meanwhile, in musicians, the corticomotor excitability and the FDI motor maps remained stable across days. In musicians, the corticomotor responses (map volume, MEP amplitude, and the rMTs) were linked to the weekly and accumulated training, indicating that those changes are associated with sensorimotor experience and related to use-dependent plasticity. In terms of the relationship between corticomotor excitability and pain sensitivity, notably, musicians reported lower NGF-related pain intensity than non-musicians in this cohort [48]. Altogether, these findings demonstrate that preexisting and established use-dependent plasticity in the motor cortex may counteract the modulatory effects of prolonged musculoskeletal pain.

### Corticomotor Changes During Prolonged Pain

Previous research has highlighted the divergent patterns of corticomotor map changes in response to pain, with some individuals showing decreases while others show facilitation [40,43]. This dichotomy in corticomotor responses has been attributed to (i) pain intensity, where individuals who show a higher decrease in corticomotor map volume also report higher pain intensity during prolonged pain conditions [9,40], and (ii) motor adaptations, where individuals who display a higher decrease in corticomotor excitability also display reduced variability in motor performance [43]. In this sense, a recent cross-sectional study indicated that musicians with chronic pain did not exhibit significantly different levels of corticomotor excitability and motor maps than pain-free musicians, suggesting that long-term motor training could counteract the effects of chronic pain on the motor system [47]. The current findings not only confirm and expand those previous results but also underscore that the interindividual variability in pain intensity responses, and its relationship with corticomotor excitability, is associated with the prior motor experience.

Musical training, characterised by repetitive and precise motor actions, induces use-dependent plasticity, leading to corticomotor adaptations and refined motor maps [18,22,26,30,47]. Use-dependent plasticity is a neurobiological phenomenon that contributes to the shape and strength of connections in the motor system [5]. Robustness and consolidation of use-dependent plasticity have been interpreted as the result of Hebbian activity-dependent synaptic plasticity [17]. Essentially, this means that synapses engaged in neuronal pathways activated during specific tasks or activities become potentiated over repetition and time, resulting in long-term plasticity [11,44]. Following this notion, it is likely that the finding of no significant effect of prolonged pain on corticomotor excitability in musicians can be explained by the strong, long-term use-dependent plasticity acquired through years of intensive and repetitive training of precise fine-motor actions [18]. Pre-existing and robust synaptic plasticity, improved through repetitive and precise motor actions, might modulate the trajectory of maladaptive pain plasticity, potentially mitigating its effects. This training-induced priming could stabilize the effects of adversarial plasticity mechanisms associated with pain [47,50,53], possibly by maintaining corticomotor excitability at a level where further reductions are less likely to occur.

Alternatively, another mechanism could relate to the motor and pain coping strategies developed by musicians compared to their peers’ non-musicians. Musicians are disproportionately affected by chronic pain compared to non-musicians [24,52]. However, despite the high prevalence of pain among musicians, they often continue to perform, raising even questions about the impact of pain on musicians’ well-being and performance [42]. Previous evidence indicates that, compared to controls, trained musicians show decreased activity and functional connectivity of the insula, a key region implicated in pain processing, in response to sensory perturbations affecting their performance [21,23] and during chronic pain conditions [50]. Furthermore, musicians experiencing chronic pain report fewer interferences with their daily activities compared to non-musicians with chronic pain and maintain their practice levels comparable to pain-free peers [47,50]. These findings suggest that individuals who perform extensive motor training under high psychological and physical demands, like musicians, might develop coping strategies that reduce the impact of pain on performance. In this study, musicians rated their pain as less intense than non-musicians and exhibited a relationship between higher weekly musical training and both lower pain scores [48] and smaller motor map volumes. It is conceivable, therefore, that musicians’ coping mechanisms, such as dissociating movements from sensory distress, might facilitate modulating pain signalling and minimising its impact on corticomotor excitability, allowing them to sustain regular training. This idea aligns with prior experiments investigating the relationship between motor performance and corticomotor excitability during prolonged experimental pain. Research showed that individuals with higher motor variability in the presence of prolonged pain also exhibit higher corticomotor excitability [43]. Moreover, other motor approaches, like action observation coupled with motor imagery, have shown evidence for counteracting corticomotor depression associated with pain [25]. Thus, an interpretation of our results could be that different motor strategies during prolonged pain may actively influence the effects of pain on corticomotor excitability. However, this detailed mechanism was not studied in this study, so further investigation is necessary.

### Limitations

While we controlled the effects of induced pain on hand and arm functions through qualitative assessment, we did not account for other quantitative activities, such as the number of training hours, particularly among musicians. Additionally, our study did not control for objective measures of motor function or coping strategies, which could have shed light on whether the observed differences in corticomotor excitability are linked to varied movement patterns. To address these limitations, future studies should incorporate monitoring daily and physical activity levels, along with more precise measures of motor and psychological performance, to provide a clearer understanding of the relationship between movement, corticomotor excitability, and pain processing.

## Conclusion

This study revealed that in musicians, a model to investigate use-dependent corticomotor plasticity, experimentally-induced prolonged pain showed a non-significant effect on the corticomotor excitability and elicited lower pain intensity than in non-musicians where decreased motor maps were found. These novel results reinforce the notion that prior motor training and its associated use-dependent plasticity may counteract the effects of persistent pain in the motor system. The dissociation of pain responses as a function of prior use-dependent plasticity may represent a potential factor to phenotype patients with chronic pain.

